# Identification of Adjuvants that Enhance the Magnitude and Durability of Antibody Responses to a Nanoparticle-Based Malaria Vaccine

**DOI:** 10.64898/2025.12.05.692661

**Authors:** Yogesh Nepal, Alexandra Francian, Yevel Flores-Garcia, Bryce T. Roberts, Rabia Khan, Garrett Wondra, Aurélie Demierre, Falko Apel, Fidel Zavala, Bryce Chackerian

## Abstract

To evaluate strategies for increasing vaccine efficacy, we assessed the ability of different classes of adjuvants to enhance vaccine-induced antibody responses to a Qβ bacteriophage virus-like particle (VLP)-based vaccine for malaria targeting the vulnerable L9 epitope from the circumsporozoite protein (CSP). From a panel of over a dozen adjuvants, five formulations significantly increased anti-CSP IgG titers and two also enhanced antibody durability. When assessed in a mouse malaria challenge model, immunization of mice with Qβ L9 VLPs plus either of two promising adjuvants, Cquim-MA and LQ, elicited sterilizing protection when mice were challenged soon after vaccination and also six months later. This study identifies promising adjuvants with different formulations that enhance both the magnitude of CSP-targeting antibodies and the longevity of VLP-induced immunity.

**IMPORTANCE:** Effective pre-erythrocytic *Plasmodium falciparum* vaccines targeting CSP must elicit strong and durable antibody responses. Current malaria vaccines are effective soon after immunization, but their efficacy decreases as antibody titers wane. Here, we evaluated the compatibility of a diverse panel of novel adjuvants in combination with a nanoparticle-based vaccine that targets an epitope within CSP that is particularly vulnerable to antibodies. Two adjuvants, with different formulations and mechanisms of action, elicited strong and durable antibody responses in mice and could elicit short-and long-term sterilizing immunity from malaria infection.

## INTRODUCTION

*Plasmodium falciparum* (*Pf*) malaria is a devastating disease responsible for over 250 million infections and over 600,000 deaths in 2024 (1). Vaccines targeting multiple stages of the *Pf* lifecycle are in development, but targeting pre-erythrocytic parasite offers the opportunity to completely prevent infection by neutralizing infectious sporozoites (2). However, sporozoites are able to reach the liver within 1-3 hours of inoculation (3, 4), meaning that there is only a brief period after exposure to a *Pf*-infected mosquito during which antibodies can effectively target the parasite (5). Furthermore, productive blood-stage infection can be seeded by just a single infected hepatocyte (6). Thus, the efficacy of sporozoite-targeted vaccines, especially vaccines that aim to induce liver infection-blocking antibodies, is dependent on the ability to elicit both strong and exceptionally durable immune responses (7, 8).

Two pre-erythrocytic vaccines that target the *Pf* circumsporozoite protein (CSP), the most abundant protein on the surface of sporozoites, are approved for clinical use in malaria-endemic regions. RTS,S/AS01_E_ and R21/Matrix-M provide protection from infection, but their efficacy wanes over time as anti-CSP antibody levels decline (9-12). The limitations of these vaccines have prompted the search for new strategies that yield stronger and more durable protection, including targeting more vulnerable, less immunodominant, epitopes within CSP (13). While current approved vaccines target the immunodominant major repeat region of CSP, a promising alternative target is the minor repeat CSP epitope targeted by the L9 monoclonal antibody (mAb) (14). Passive administration with L9 mAb provides sterilizing immunity in mouse infection models at lower doses than other mAbs that target immunodominant major repeat epitopes (14). Moreover, passive immunization with L9 mAb provides protection from infection in experimental human challenge studies and field trials in malaria-endemic areas (15, 16). We produced a vaccine targeting the L9 epitope by displaying a peptide representing this epitope at high density on the surface of an immunostimulatory vaccine platform derived from Qβ bacteriophage virus-like particles (VLPs) (17, 18). In mice, immunization with Qβ L9 VLPs elicited high-titer and durable anti-CSP antibody responses that significantly reduced parasite liver burden in a mouse malaria challenge model (17, 18). Although VLP-based vaccines are highly immunogenic, the use of adjuvants to increase anti-CSP antibody levels was critical for the induction of sterilizing immunity (17, 18). We identified Advax-3, a TLR9 agonist (CpG55.2) combined with aluminum hydroxide, as an adjuvant which increased anti-CSP antibody levels by 8-fold (19), leading to sterile protection (i.e. an absence of blood-stage parasites) in 60% of challenged mice (17). However, use of Advax-3 only transiently increased titers; within 2-3 months after immunization, the levels of anti-CSP antibodies were comparable to mice immunized without exogenous adjuvant (19). These results highlight the need for alternative adjuvants that elicit more durable antibody responses. Here, we report the evaluation of a selection of nine adjuvants with different formulations and mechanisms of action for their ability to increase the magnitude and durability of anti-CSP antibody responses elicited by Qβ L9 VLPs. Two promising adjuvants were identified and evaluated for short-and long-term efficacy in malaria challenge studies in a mouse model.

## RESULTS

### Evaluating the compatibility of diverse adjuvants with Qβ L9 VLPs

We initially screened 20 distinct adjuvant formulations comprising various TLR agonists, liposomes, emulsions, and other proprietary compounds developed by different academic and commercial groups and obtained through the NIAID adjuvant development program(20). This manuscript reports the immunogenicity of a subset of nine adjuvants that were selected to represent diverse mechanistic classes and formulations (**Table 1**). The adjuvants included in this study are formulated as liposomes (LQ, LMQ, Lipo2B182C, Lipo2B182C+1V270), contain QS-21 or synthetic saponins (LQ, LMQ, TQL-1055), and/or contain TLR4 agonists (LMQ, Lipo2B182C, and Lipo2B182C+1V270). We also included one squalene oil-in-water emulsion formulation (Sepivac SWE™; hereafter called SWE)) and several adjuvants which contain small molecule TLR7/8 agonists (Cquim-MA, AHQ-II, Auroxiquim, and Lipo2B182C+1V270).

**Table 1.**
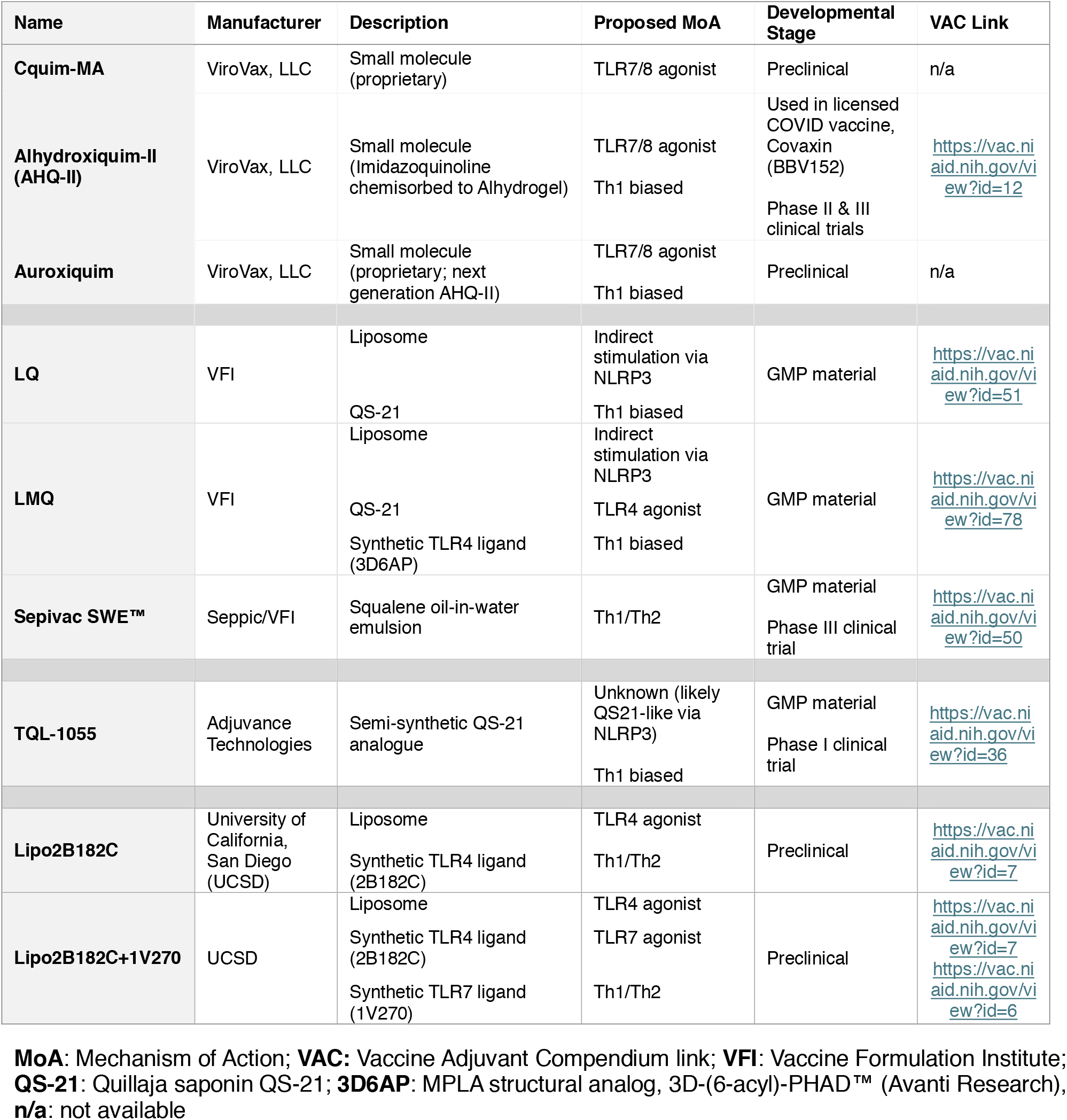
Adjuvant characteristics. Summary of adjuvants used in this study, including manufacturer, formulation type, proposed mechanism of action (MoA), and developmental stage. More information can be found at the NIAID Vaccine Adjuvant Compendium (VAC) entries cited within the table.

To evaluate immunogenicity, each adjuvant was combined with a standard 5 µg dose of Qβ L9 VLPs and was administered intramuscularly to female BALB/c mice three times at three-week intervals. Sera were collected at two time points; we measured peak antibody levels, (three weeks following the final immunization), and the longevity of antibody responses (6-7 months following the final immunization) (**Fig. 1a**). Anti-CSP IgG titers were measured by ELISA against full-length recombinant CSP and compared to mice immunized with Qβ L9 VLPs without exogenous adjuvant.

**Fig 1.**
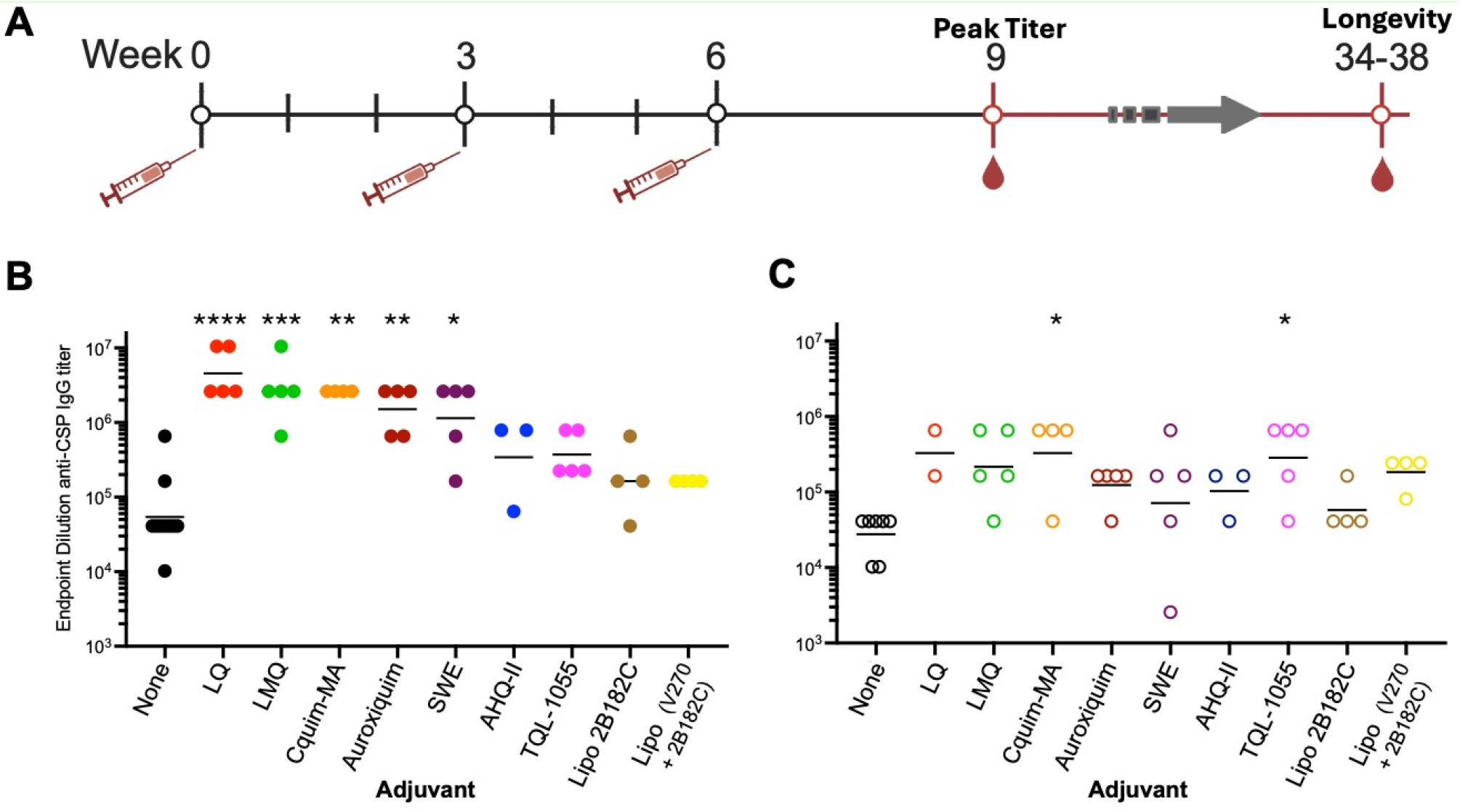
Effects of adjuvants on the anti-CSP IgG titers elicited by Qβ L9 VLPs. **a** Schematic of the immunization schedule. Female BALB/c mice were immunized intramuscularly at weeks 0, 3, and 6 with 5 µg of Qβ L9 VLPs formulated with or without adjuvant. Sera were collected 3 weeks following the third immunization at week 9 and ∼6 months later, between weeks 34 and 38. **b** Anti-CSP IgG titers measured by ELISA against full-length recombinant CSP using sera collected at week 9 post immunization or **c** collected at 34 weeks (unadjuvanted, Cquim-MA, TQL-1055, Lipo 2B182C, and Alhydroxyquim-II) or 38 weeks (LQ, LMQ, and SWE) following the final immunization. Each dot represents an individual mouse. Horizontal bars indicate geometric means, and error bars represent standard error of the mean (SEM). Statistical comparisons between adjuvanted groups and the unadjuvanted group was performed using Kruskal-Wallis with Dunn’s post-hoc test; * denotes p < 0.05; **, p < 0.005; ***, p < 0.0005, ****, p < 0.0001.

Immunization with Qβ L9 VLPs alone (unadjuvanted control group) elicited high anti-CSP IgG titers, reflecting the ability of peptide-conjugated VLPs to elicit strong antibody responses due to the multivalent display of the target epitope (21, 22) and potentially because Qβ VLPs naturally encapsidate bacterial-derived single-stranded RNA (a TLR7/8 agonist), which modestly enhances their potency (23). However, combining this vaccine with each of the adjuvant formulations reported here resulted in higher geometric mean anti-CSP IgG titers compared to unadjuvanted Qβ L9 VLPs; five of the nine adjuvant formulations significantly increased antibody titers (**Fig. 1b**). LQ (24), a liposomal formulation containing the saponin QS-21, resulted in an 84-fold increase in anti-CSP IgG titers, LMQ (liposomes containing QS-21 and a synthetic TLR4 agonist) (24, 25) and Cquim-MA (dual TLR7/8 agonist) (26) resulted in a 49-fold increase in titers, Auroxiquim (dual TLR7/8 agonist) resulted in a 28-fold increase, and SWE (squalene oil-in-water emulsion) (27) resulted in a 21-fold increase. TQL-1055(28), Alhydroxyquim-II (29), and Lipo2B182C or Lipo2B182C+1V270 (30) increased anti-CSP antibody titers, but failed to reach statistical significance compared to Qβ L9 VLPs alone.

To assess whether the adjuvants evaluated in this study elicit sustained elevated antibody titers, anti-CSP IgG titers were also measured >6 months following the final immunization (**Fig. 1c**). Although antibody levels declined in nearly all of the adjuvanted groups over time, nearly all of the mice which received adjuvanted Qβ L9 VLPs formulations had higher antibody levels than mice that received unadjuvanted Qβ L9 VLPs. Mice immunized with Qβ L9 VLPs adjuvanted with LQ, TQL-1055 and Cquim-MA elicited the highest antibody titers at this timepoint, with a ∼12-fold increase in titers compared to unadjuvanted Qβ L9 VLPs. Use of the six other adjuvants also trended towards higher antibody titers than the unadjuvanted control at this timepoint, ranging from 2-to 8-fold higher than the control group, but these increases were not significantly different.

### Effects of adjuvants on antibody avidity

To further evaluate the effects of adjuvants on antibody responses, we measured the avidity of anti-CSP antibody responses and the levels of specific IgG isotypes. Clinical efficacy of RTS,S/AS01_E_ has been correlated with both higher antibody titer and increased avidity (31, 32), but it is unclear whether antibody avidity affects the functionality of antibodies that target the L9 epitope. To evaluate the effects of different adjuvants on the binding strength of antibodies generated against CSP, we compared antibody avidities amongst the groups using serum samples that were obtained at peak antibody titers using a chaotrope-based avidity assay. None of the adjuvanted groups elicited statistically significant changes in avidity relative to non-aduvanted vaccines, although there was a trend toward higher avidities in the groups with lower antibody titers (**Fig. 2**).

**Fig 2.**
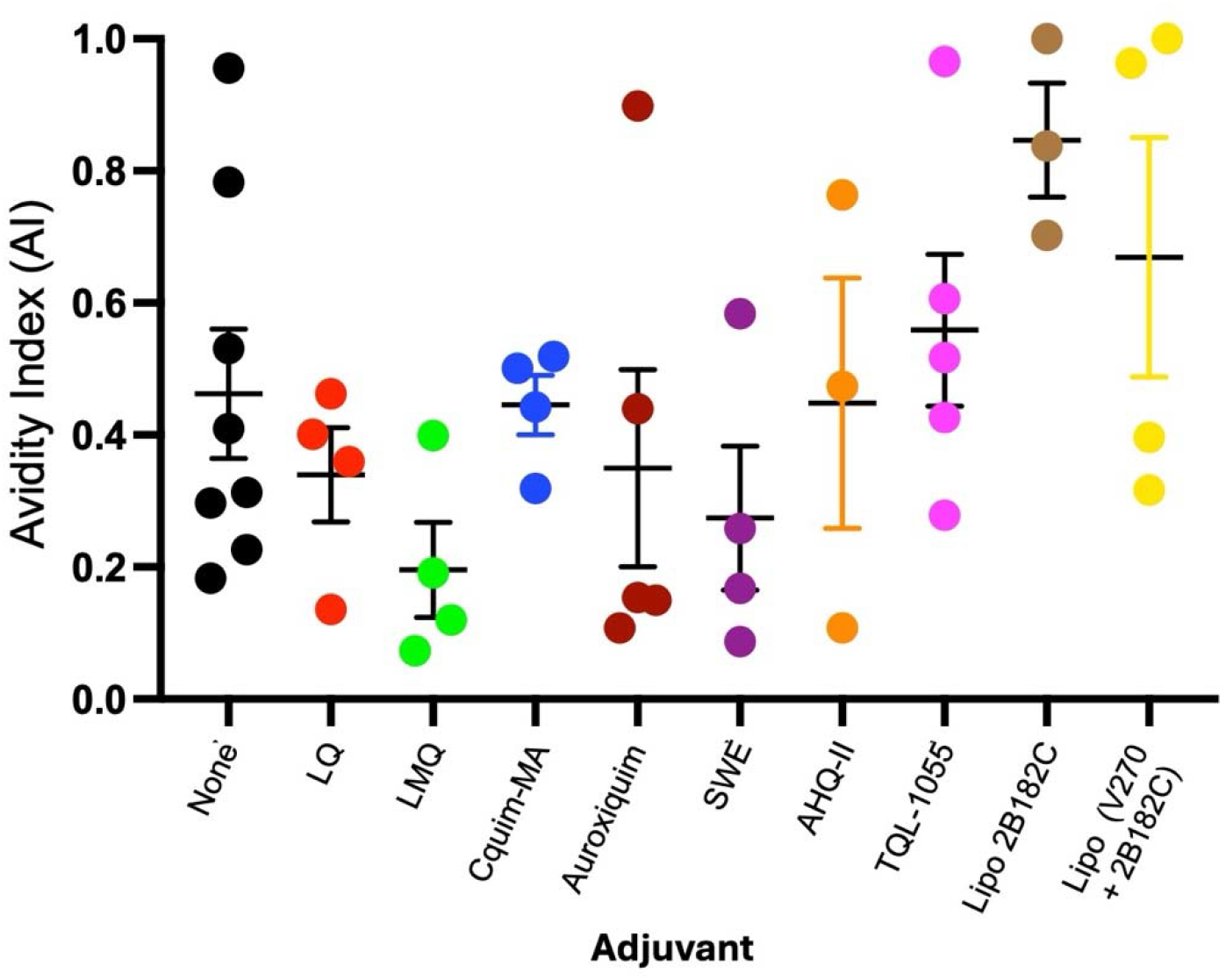
Adjuvants influence the avidity of the anti-CSP IgG response. Avidity was measured by urea-based ELISA using sera collected at peak titers. Avidity index (AI) was calculated as the ratio of absorbance in 8M urea-treated wells to that in PBS-treated wells (AI = A_450_Urea/A_450_PBS. Each point represents the AI for an individual mouse. Bars indicate group means; error bars represent SEM.

### Adjuvants skew IgG isotypes

To assess isotype skewing by different adjuvants, we measured CSP-specific IgG1 antibodies (associated with Th2 responses) and IgG2a antibodies (associated with Th1 responses) induced by Qβ L9 VLPs in combination with the different adjuvants. **Fig. S1** shows the log_10_-transformed ratio of IgG2a to IgG1 for each group of immunized mice. On average, unadjuvanted Qβ L9 VLPs induced balanced IgG2a:IgG1 responses, although there was considerable variability in this ratio in individual mice. Most of the adjuvants that were evaluated elicited antibody responses that were slightly biased towards IgG2a, suggesting that these adjuvants largely induced Th1 responses. Lipo2B182C and Lipo2B182C+1V270 elicited balanced responses, and Auroxiquim skewed responses towards IgG1.

### Qβ L9 VLPs plus selected adjuvants provide long-lasting protection from malaria infection

We hypothesized that the ability of adjuvants to promote durable antibody responses would result in sustained sterilizing protection from malaria infection. To evaluate this question, groups of C57BL/6 mice were vaccinated with Qβ L9 VLPs in combination with Cquim-MA or LQ. Cquim-MA and LQ were selected because of their ability to enhance the magnitude and longevity of anti-CSP antibodies. Following immunization, mice were challenged with mosquitoes carrying transgenic *Plasmodium berghei* (*Pb*) sporozoites expressing full-length *Pf*CSP and luciferase (*Pb*-*Pf*CSP-luc) (33) at three weeks after the final immunization, in order to assess protection when antibody levels were at their highest, or at six months following the final immunization, to assess the durability of both antibody levels and protection.

We have previously shown that immunization with Qβ L9 VLPs in combination with Cquim-MA can elicit sterilizing immunity in ∼40-80% of vaccinated mice that were challenged soon after vaccination (18). To confirm that Qβ L9 VLPs adjuvanted with LQ could provide similar levels of sterilizing immunity, we repeated this study in groups of mice immunized with three doses of Qβ L9 VLPs plus Cquim-MA or LQ. Two weeks following the final dose, anti-CSP antibody concentrations were measured using a quantitative ELISA that utilizes a standard curve generated using known concentrations of the anti-CSP monoclonal antibody 2A10 (34). Mice that received vaccine plus Cquim-MA or LQ showed similar mean antibody concentrations (**Fig. 3a**). Three weeks following the final dose, mice were challenged with mosquitoes carrying *Pb*-*Pf*CSP-luc sporozoites; infection was monitored by evaluating liver parasitemia ∼two days post-infection, as measured by liver luminescence, and blood-stage parasitemia beginning at day four post-infection, as measured by daily Giemsa staining of blood smears. Compared to unvaccinated mice, Qβ L9 VLP vaccinated mice, combined with either adjuvant, had low liver parasite loads (**Fig. 3b**) that were only slightly higher than the background luminescence levels observed in naïve, noninfected (NNI) mice. There was no significant difference in liver loads in mice that received vaccine plus Cquim-MA versus vaccine plus LQ. In both groups, 4 of 7 mice (57%) were completely protected from blood-stage malaria infection (**Fig. 3c**). Thus, both Cquim-MA and LQ adjuvants combined with Qβ L9 VLPs provide similarly strong protection from malaria infection.

**Fig 3.**
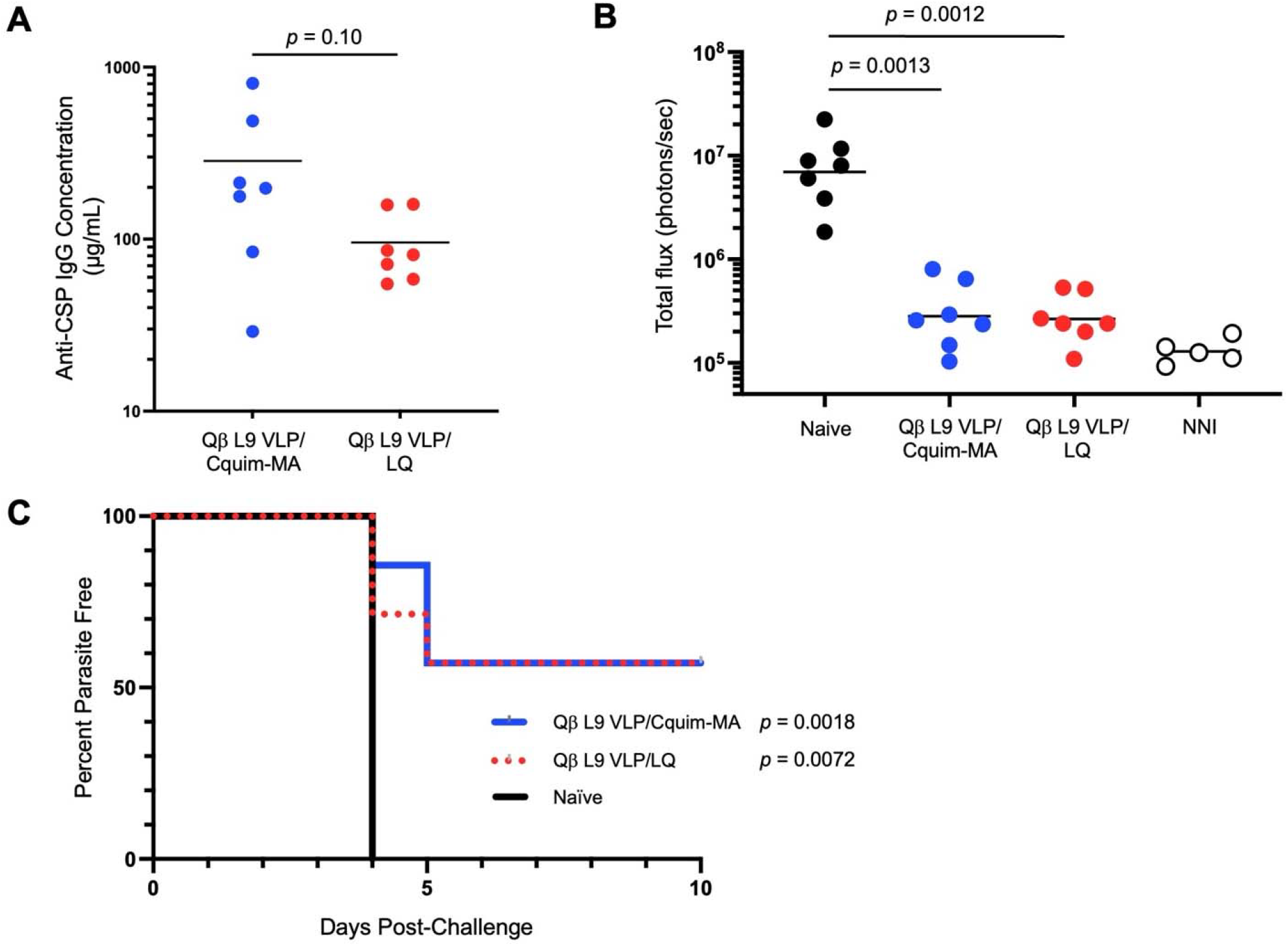
Immunization with Qβ L9 VLPs plus adjuvants elicits strong antibody responses and protects against malaria infection. C57BL/6 mice were immunized three times at weeks 0, 3, and 6, followed by a challenge with five *Pb-PfCSP-Luc*-infected mosquitoes at week 9. Liver luminescence was measured 42□h after the mosquito challenge, and blood smears were taken starting on day 3 post-infection. **a** Anti-CSP antibody levels in serum following the third vaccination, prior to challenge. Horizontal lines represent the mean anti-CSP antibody concentration for each group. Groups are compared by 2-tailed unpaired *t* test. **b** Parasite liver burden measured via luminescence. Horizontal lines indicate geometric mean luminescence for each group. Statistical comparisons were performed using a one-way ANOVA in which experimental groups were compared to every other challenged group using Dunnett’s multiple comparisons test. NNI refers to naïve (non-vaccinated), non-infected mice. **c** Protection from blood-stage infection, as measured by percent of blood parasite-free mice post-challenge. A log-rank test was used to statistically compare vaccinated groups to naïve, infected mice.

To evaluate the durability of protection, additional groups of mice were vaccinated with Qβ L9 VLPs plus Cquim-MA or LQ and then challenged 6 months after the final dose of vaccine. To assess the durability of antibody responses, anti-CSP antibody concentrations were measured at early (2 weeks) and late (6 months) timepoints following the third vaccination. At the early timepoint, we observed similar anti-CSP IgG concentrations in the two adjuvanted groups (**Fig. 4a**); these levels are similar to what is reported in our previous challenge experiment (**Fig. 3a**). By six months after vaccination, mean anti-CSP antibody levels had decreased by ∼4-fold in both groups (**Fig. 4b**). Nevertheless, these antibody levels were still high; mean concentrations were 53 µg/mL in the group that received LQ and 42 µg/mL in the group that received Cquim-MA. Upon mosquito challenge, both groups had reduced liver loads relative to unvaccinated mice (**Fig. 4c**), although this reduction was less pronounced compared to our previous challenge (**Fig. 3**) where mice were challenged early after vaccination. Blood-stage parasite were detected in all seven unvaccinated mice by day 4, whereas two of seven mice in both of the vaccinated groups were protected from infection (**Fig. 4d**). Thus, even six months after vaccination, a subset of mice were completely protected from malaria infection.

**Fig 4.**
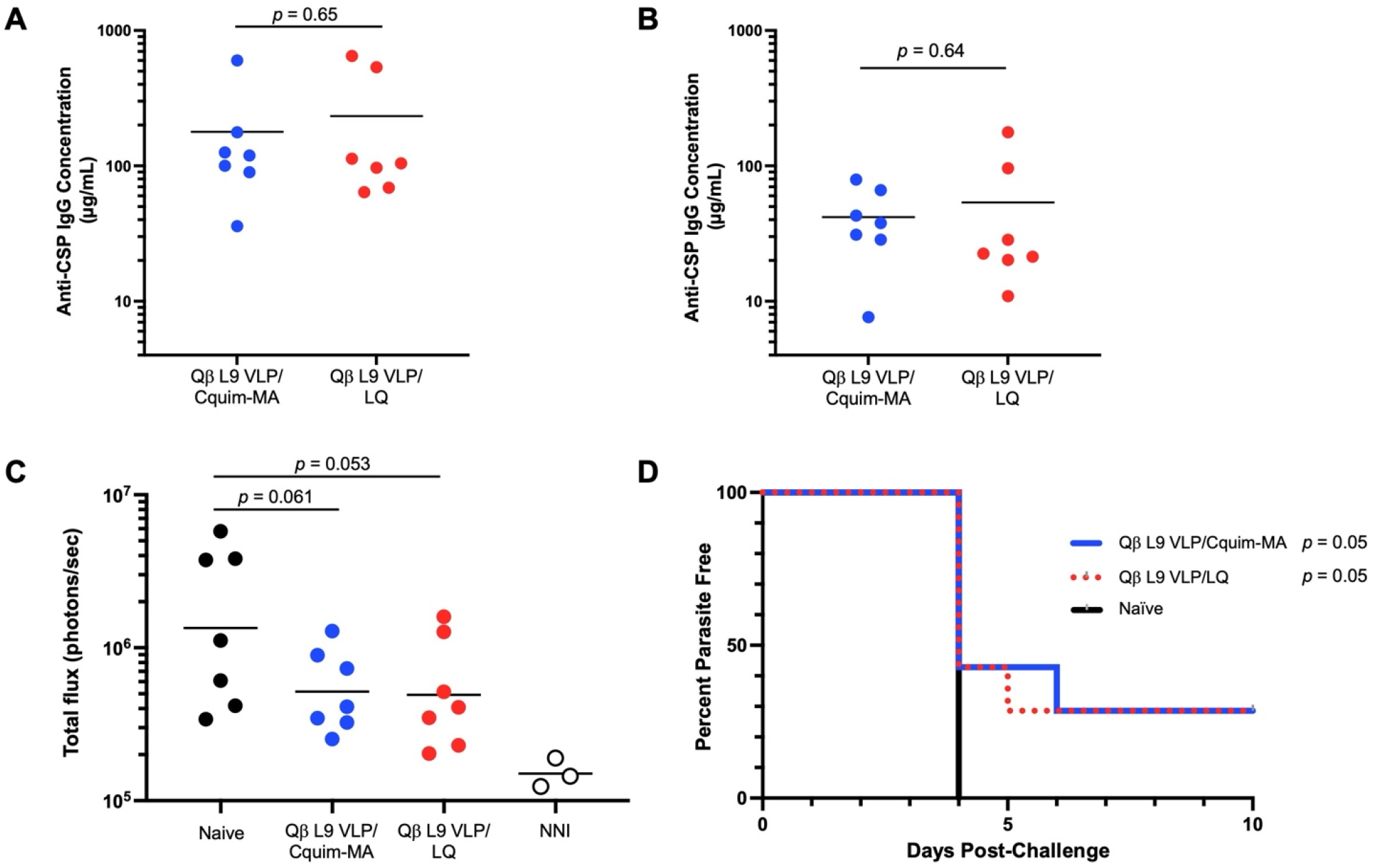
Immunization with Qβ L9 VLPs plus adjuvants provides durable protection from malaria infection. C57BL/6 mice were immunized three times at weeks 0, 3, and 6. Sera was collected at week 8 and week 32. Mice were challenged with five *Pb-PfCSP-Luc*-infected mosquitoes at week 34. Liver luminescence was measured 42□h after the mosquito challenge, and blood smears were taken starting on day 3 post-infection. Anti-CSP antibody levels were measured in serum obtained prior to challenge, at (**a**) week 8 or (**b**) week 32. Horizontal lines represent the mean anti-CSP antibody concentration for each group. Groups are compared by 2-tailed unpaired *t* test. **c** Parasite liver burden measured via luminescence. Horizontal lines indicate geometric mean luminescence for each group. Statistical comparisons were performed using a one-way ANOVA in which experimental groups (adjusted by subtracting background luminescence) were compared to every other challenged group using Dunnett’s multiple comparisons test. NNI refers to naïve (non-vaccinated), non-infected mice. **d** Protection from blood-stage infection, as measured by percent of blood parasite-free mice post-challenge. A log-rank test was used to statistically compare vaccinated groups to naïve, infected mice.

## DISCUSSION

In this study, we paired a promising epitope-targeted malaria vaccine with a panel of emerging adjuvants possessing distinct immunostimulatory properties and evaluated how each adjuvant affected peak anti-CSP IgG antibody titers and the durability of antibody responses. Our results identified several promising adjuvants that substantially increased anti-CSP antibody levels over an extended period following immunization. Liposome-based adjuvants containing QS-21 (LQ and LMQ) outperformed most other adjuvants in regard to peak titers, but surprisingly, the addition of a synthetic TLR4 agonist to this formulation (LMQ) did not significantly increase antibody titer nor increase durability, although this may be related to the varying saponin dosages in these adjuvants. Another promising adjuvant, Cquim-MA (TLR7/8 agonist), does not incorporate either TLR4 agonists or saponins, but was able to elicit comparably high titers and durability as LQ. These adjuvants leverage varying innate sensing mechanisms, mechanisms of action, and immune activation profiles, yet they ultimately resulted in similarly high IgG titers and durability. This illustrates the importance in using an empirical adjuvant selection process on a vaccine-specific basis (35).

Adjuvants affect other features of antibodies, such as avidity and isotype, which may impact vaccine efficacy. In this study, we show that in Balb/c mice, different adjuvants elicit different ratios of IgG2a:IgG1, which is reflective of an ability to elicit Th1 versus Th2 responses. Although there is some evidence that specific IgG isotypes may correlate with malaria protection during natural infection (36), a recent study showed that anti-CSP monoclonal antibodies which lack a functional Fc region retain protective efficacy *in vivo* (37). These data suggest that it is unlikely that differences in the IgG isotypes elicited by L9 VLPs will affect protective efficacy against malaria. Nevertheless, understanding how adjuvants influence antibody isotype may be important for other VLP-based vaccines that are under development.

Interestingly, our data indicated that there was a trend indicating an inverse correlation between antibody titer and avidity in the adjuvanted groups. It is possible that adjuvants that allow for the induction of higher antibody levels do so by promoting the expansion of B cells with lower affinity B cell receptors, potentially through an extrafollicular pathway (38). Multivalent particle-based vaccines have been shown to preferentially allow low-affinity B cells to enter germinal centers and differentiate into plasmablasts (39). Future studies that explore the dynamics of how adjuvants combine with VLP-based immunogens to generate both short-lived and long-lived plasmablasts could shed light on how these different mechanisms interact and shape the affinity and durability of antibody responses.

We have previously demonstrated that Cquim-MA can enhance the ability of Qβ L9 VLPs to reduce parasite liver burden and provide sterilizing immunity in a mouse malaria challenge (26). This work identifies a second adjuvant (LQ) with a similar ability to enhance anti-CSP antibody responses and provide strong protection from malaria infection in a mouse malaria challenge model. We also assessed the durability of protection by following mice for 6 months post-immunization and measuring antibody concentrations and protective efficacy. Although anti-CSP antibody levels declined an average of 4-fold, we still observed sterilizing protection in nearly 30% of immunized mice. Thus, adjuvanted L9 VLPs can provide durable, protective responses in mice for up to 6 months post-vaccination. Although it is unclear whether efficacy will be maintained after 6 months, we previously showed that anti-CSP antibody levels in mice immunized with VLP-based vaccines targeting the L9 epitope (in combination with Cquim-MA) drop in the weeks following immunization, as short-lived plasmablasts decline, but then stabilize (18). In contrast, pediatric trials of RTS,S/AS01_E_ have shown that this vaccine largely elicits short-lived anti-CSP antibody responses, and that antibody levels and vaccine efficacy steadily decline after immunization (40). Thus, use of an adjuvanted VLP-based vaccine targeted a vulnerable epitope in CSP may represent one strategy for prolonging antibody levels and vaccine efficacy.

## METHODS

### Production of Qβ L9 VLPs

Qβ bacteriophage VLPs were produced and purified by expression in *Escherichia coli*. Qβ bacteriophage coat protein was overexpressed from plasmid pETQCT using electrocompetent *E. coli* C41 (DE3) cells (Sigma-Aldrich)(19). Bacterial pellets containing self-assembled VLPs were resuspended in lysis buffer (100 mM NaCl, 10 mM EDTA, 50 mM Tris-HCl, and 0.45% deoxycholate) and incubated on ice for 30 min, followed by three to five cycles of sonication at 20 Hz, until the solution was clear. Following sonication, residual DNA was removed using 10 µg/mL DNase, 2.5 mM MgCl_2_, and 0.05 mM CaCl_2_ (all final concentrations) upon incubation on ice for 1 h. The supernatant was isolated by centrifugation at 10,000 rpm for 30 min (TA-14-50 fixed-angle rotor). Protein was precipitated by adding ammonium sulfate to the supernatant to make a 70% solution, followed by incubation on ice for 15 min. Precipitated protein was spun at 10,000 rpm for 15 min, and the pellet was resuspended in cold SCB buffer (10 mM Tris-HCl, 100 mM NaCl, and 0.1 mM MgSO_4_) and then subjected to size exclusion chromatography using Sepharose CL-4B beads (Sigma). Fractions that contained Qβ VLPs were identified by agarose gel electrophoresis, incubated in 70% ammonium sulfate overnight at 4°C to precipitate protein, and then pelleted by centrifugation at 10,000 rpm for 15 min. VLPs were resuspended in phosphate-buffered saline (PBS) (pH 7.4) and dialyzed two times against PBS (pH 7.4). Prior to peptide modification, Qβ VLPs were depleted of endotoxin by four rounds of phase separation using Triton X-114 (Sigma-Aldrich) (41). The final concentration of Qβ VLPs was determined via SDS-PAGE using known concentrations of hen egg lysozyme as standards.

Qβ VLPs were chemically conjugated to a synthetic peptide corresponding to the L9 epitope of CSP. The L9 peptide containing a C-terminal glycine-glycine-glycine-cysteine linker (NANPNVDPNANPNVD-GGGC) was synthesized by GenScript. Conjugation was performed using the heterobifunctional crosslinker succinimidyl 6-[(β-maleimidopropionamido) hexanoate] (SMPH; Thermo Fisher Scientific). Qβ VLPs were first incubated with SMPH at a 10:1 molar ratio (SMPH:Qβ coat protein) for 1 h at room temperature. Excess SMPH was removed by ultrafiltration using an Amicon Ultra-4 centrifugal filter unit with a 100 kDa molecular weight cutoff (Millipore Sigma). The SMPH conjugated VLPs were then incubated with the L9 peptide at a 10:1 molar ratio (peptide:Qβ coat protein) overnight at 4°C. The peptide was covalently attached to surface-exposed lysines on Qβ bacteriophage VLPs using SMPH, as previously described (42). Conjugation efficiency was evaluated by gel electrophoresis using a 10% NuPAGE gel (Invitrogen) by quantifying the percentage of Qβ coat protein with increase molecular mass due to peptide conjugation.

### Ethics Statement for Animal Studies

All animal research was approved by the Institutional Animal Care and Use Committee at the University of New Mexico Health Sciences Center under the protocol number 22-201289-HSC and at Johns Hopkins University under permit #: MO18H419.

### Mouse immunizations

To evaluate vaccine immunogenicity, groups of 4–5-week-old female BALB/c mice (n=3-10 per group; typically n=5) were obtained from Jackson Laboratory and immunized intramuscularly with 5 µg of Qβ L9 VLPs per mouse, either alone or formulated with the adjuvant Cquim-MA (2 µg/dose), alhydroxiquim-II (AHQ-II; 50 µg/dose), auroxiquim (100 µg/dose), LQ (5 µg QS21/dose), LMQ (2.5 µg QS21/dose), Sepivac SWE™ (SWE) (1 mg squalene/dose), TQL-1055 (30 µg/dose), Lipo 2B182C (200 nMoles/dose), and Lipo(V270+2B182C) (200 nMoles/dose). Mice received a prime immunization followed by booster doses at three and six weeks after the initial injection, using the same formulation and dosing. Serum samples were collected at two time points: three weeks after the third immunization (defined as peak antibody titers) and again 28-32 weeks after the final boost (defined as long-lived antibody titers), to assess the durability of antibody responses. Note that three of the mice in the group immunized with VLPs plus LQ died during the course of the study (for reasons that were unrelated to the immunization protocol). Animals were anesthetized with 2% Isoflurane (oxygen flow of 1LL/min) during all blood collections. At the conclusion of each study, mice were euthanized by terminal cardiac puncture. Animals were anesthesized by intraperitoneal administration of 0.3 mL of a mixture of ketamine (10 mg/mL) and xylazine (1 mg/mL) prior to the blood draw. Euthanasia was confirmed by cervical dislocation.

### Malaria challenge studies

Challenge studies were performed using 7-8-week–old C57BL/6 mice, which are more susceptible to malaria challenge than Balb/c mice. Groups of mice were immunized with 5 μg of VLPs with Cquim-MA adjuvant (2 µg) or LQ adjuvant (5 µg QS21/dose) on days 0, 21, and 42. Negative control mice were unimmunized. In studies to evaluate short-term efficacy, sera were collected three weeks following the final boost and then mice were challenged two days later. To evaluate long-term vaccine efficacy, sera were collected three weeks and six months following the final boost; mice were challenged two days after the final sera collection.

Challenges were performed using *Anopheles stephensi* mosquitoes infected with transgenic *P. berghei* sporozoites engineered to express luciferase and full-length *Pf*CSP in place of *P. berghei* CSP (*Pb*-*Pf*CSP-luc) (33). Infected mosquitoes were generated by allowing the insects to blood-feed on *Pb*-*Pf*CSP-luc-infected mice. For challenges, mice were anesthetized prior to exposure using 2% Avertin. Each mouse was then exposed to five infected mosquitoes for a 10 min blood meal. After feeding, the number of mosquitoes with visible blood meals was recorded. Liver luminescence was measured 42 h post-challenge. Mice were sedated using 2-4% isoflurane, and 100 μL of D-luciferin (30 mg/mL) was administered via intraperitoneal injection. Luminescence from the liver was subsequently captured using the IVIS Spectrum Imaging System (Perkin Elmer) while the mice were under sedation using isoflurane. Starting 4 days post-challenge, blood smears were collected daily and stained with Giemsa to monitor parasitemia.

### Measurement of antibody titers

Antibody titers against full-length CSP were measured by ELISA using recombinant CSP expressed in *Pseudomonas fluorescens* (43), provided by Gabriel Gutierrez (Leidos, Inc.). Immulon 2 HB ELISA plates (ThermoFisher Scientific) were coated with 200 ng of CSP in 50 µL PBS per well and incubated either overnight at 4°C or for 2 h at room temperature. Plates were then blocked with 100 µL of PBS containing 0.5% milk for 1-2 h at room temperature. Serum samples from immunized mice were serially diluted in PBS with 0.5% milk, added to the plates, and incubated for 2 h at room temperature. CSP-specific antibodies were detected using horseradish peroxidase (HRP)-conjugated goat anti-mouse IgG (Jackson ImmunoResearch, 1:4000 dilution in PBS with 0.5% milk), and signal was developed using TMB substrate and reaction was stopped using 1% HCl. Endpoint titers were defined as the highest serum dilution yielding an OD_450_ value at least two-fold above background. For challenge experiments, CSP-specific antibody concentrations in mouse sera were quantified by linear regression analysis using a standard curve generated using known concentrations of the anti-CSP central repeat-reactive mouse monoclonal antibody 2A10, as described previously (17, 19).

### Measurement of antibody isotypes

To account for variations in antibody concentrations across groups, serum dilutions yielding comparable OD_450_ values (∼2.0) for the total IgG ELISA (described above) were selected for analysis. CSP-specific IgG subclasses were measured using the same ELISA protocol as described above, substituting the secondary antibody with HRP-conjugated goat anti-mouse IgG1 or IgG2a (Jackson ImmunoResearch, 1:5000 dilution in PBS with 0.5% milk).

### Measurement of antibody avidity

Anti-CSP antibody avidity was evaluated using an ELISA-based chaotrope avidity assay (19). This assay follows the standard ELISA protocol outlined above, with the modification that after incubation with mouse serum, wells were treated with either PBS or 8 M urea, to disrupt low-affinity interactions, for 12 min. Serum was diluted similar to the isotype analysis, as described above. The avidity index (AI) was calculated as the ratio of absorbance in urea-treated wells (A_8M UREA_) to that in PBS-treated control wells (A_PBS_), expressed as: AI = A_8M UREA_ / A_PBS_. A single dilution per sample was analyzed, and all measurements were performed in triplicate.

### Statistical analysis

All statistical analyses were performed using GraphPad Prism version 11.1. In **Fig. 1**, statistical significance was tested between groups of immunized animals using Kruskal-Wallis tests with Dunn’s post-hoc test to control for multiple comparisons. In **Fig. 3a, 4a**, and **4b**, significance was tested by unpaired t-test. In **Fig. 3b** and **4c**, significance was tested using a one-way ANOVA in which experimental groups were compared to every other group. In **Fig. 3c** and **4d**, a log-rank test was used to statistically compare vaccinated groups to naïve, infected mice.

## Data availability

All data generated or analysed during this study are included as data points in this published article. Raw data is available upon request.

## ACKNOWLEDGEMENTS

We thank our partners for the generous donation of adjuvants used in this study, including Sunil David (ViroVax LLC), the Vaccine Formulation Institute (VFI), Dennis Carson’s laboratory at the University of California, San Diego, and Adjuvance Technologies Inc.

## CONTRIBUTIONS

This work was conceived by B.C., F.Z., Y.N., A.F. and B.T.R. Y.N. and B.C. (all Figs.), B.T.R and R.K. (Fig. 1), G.W. (Figs. 2 and S1), and Y.F. (Figs. 3 and 4) conducted the experiments and data analysis. F.A. and A.D. contributed critical reagents to this study. A.D. also performed formulation experiments at VFI laboratories (not reported in this manuscript) and F.A. was involved in planning and reporting of these experiments. The manuscript was drafted by Y.N., A.F., and B.C.; all authors edited the manuscript and approved its submission.

## DECLARATION OF AI USE

No AI tools were used in the writing or editing of this manuscript.

