## Supplemental Figure 1 for "Identification of Adjuvants that Enhance the Magnitude and Durability of Antibody Responses to a Nanoparticle-Based Malaria Vaccine"

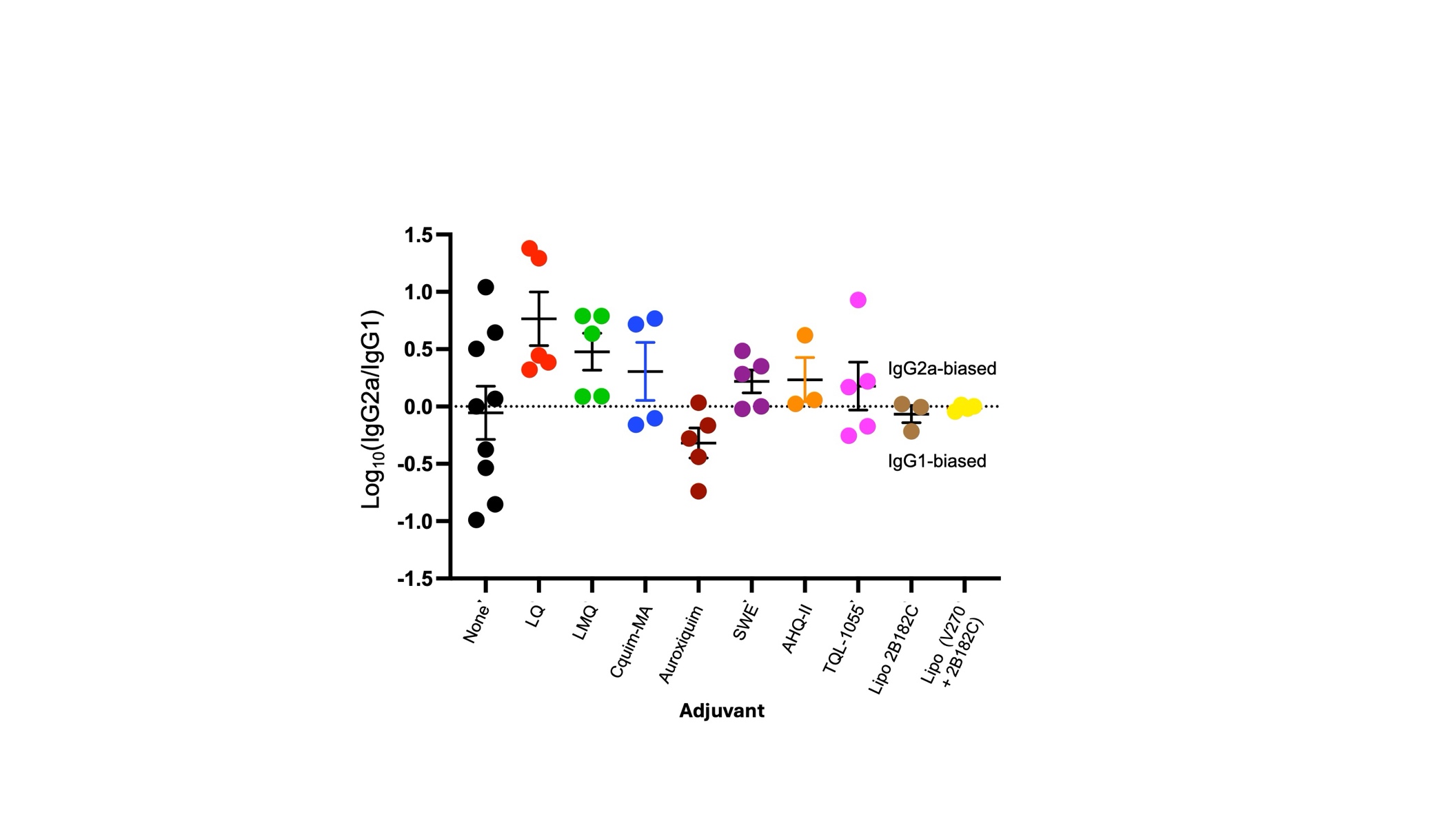

**Fig. S1. Adjuvants influence IgG isotypes.** Log₁₀-transformed ratio of CSP-specific IgG2a to IgG1 in serum collected from female BALB/c mice at peak titers. A value of 0 (dashed line) indicates a balanced IgG2a/IgG1 response. Values above or below 0 reflect IgG2a- or IgG1-skewing antibody responses, respectively. Each point represents an individual mouse; horizontal bars indicate means and error bars show SEM.
